# A derivative of the D5 monoclonal antibody that targets the gp41 N-heptad repeat of HIV-1 with broad tier-2 neutralizing activity

**DOI:** 10.1101/2020.10.23.352526

**Authors:** Adonis A. Rubio, Maria V. Filsinger Interrante, Benjamin N. Bell, Clayton L. Brown, Celia C. LaBranche, David C. Montefiori, Peter S. Kim

## Abstract

HIV-1 infection is initiated by the viral glycoprotein Env, which, after interaction with cellular coreceptors, adopts a transient conformation known as the pre-hairpin intermediate (PHI). The N-heptad repeat (NHR) is a highly conserved region of gp41 exposed in the PHI; it is the target of the FDA-approved drug enfuvirtide and of neutralizing monoclonal antibodies (mAbs). However, to date these mAbs have only been weakly effective against tier-1 HIV-1 strains, which are most sensitive to neutralizing antibodies. Here, we engineered and tested 11 IgG variants of D5, an anti-NHR mAb, by recombining previously described mutations in four of D5’s six antibody complementarity-determining regions. One variant, D5_AR, demonstrated 6-fold enhancement in ID_50_ against lentivirus pseudotyped with HXB2 Env. Importantly, D5_AR exhibited weak cross-clade neutralizing activity against a diverse set of tier-2 HIV-1 viruses, which are less sensitive to neutralizing antibodies than tier-1 viruses and are the target of current antibody-based vaccine efforts. In addition, the neutralization potency of D5_AR IgG was greatly enhanced in target cells expressing FcγRI with ID_50_ values below 0.1 μg/mL; this immunoglobulin receptor is expressed on macrophages and dendritic cells, which are implicated in the early stages of HIV-1 infection of mucosal surfaces. D5 and D5_AR have equivalent neutralization potency in IgG, Fab, and scFv formats, indicating that neutralization is not impacted by steric hindrance. Taken together, these results provide support for vaccine strategies that target the PHI by eliciting antibodies against the gp41 NHR.

**Importance:** Despite advances in anti-retroviral therapy, HIV remains a global epidemic and has claimed more than 32 million lives. Accordingly, developing an effective vaccine remains an urgent public health need. The gp41 N-heptad repeat (NHR) of the HIV-1 pre-hairpin intermediate (PHI) is highly conserved (>90%) and is inhibited by the FDA-approved drug enfuvirtide, making it an attractive vaccine target. However, to date NHR antibodies have not been potent. Here, we engineered D5_AR, a more potent variant of the anti-NHR antibody D5, and established its ability to inhibit HIV-1 strains that are more difficult to neutralize and are more representative of circulating strains (tier-2 strains). The neutralizing activity of D5_AR was greatly potentiated in cells expressing FcγRI; FcγRI is expressed on cells that are implicated at the earliest stages of sexual HIV-1 transmission. Taken together, these results bolster efforts to target the gp41 NHR and the PHI for vaccine development.

## Introduction

Over 35 million people currently live with HIV/AIDS globally (1); despite the promise of anti-retroviral therapy, a preventative HIV-1 vaccine remains an urgent global health need. Identifying and characterizing broadly neutralizing antibodies (bnAbs) is essential to informing the design of vaccine antigens that could elicit such antibodies *in vivo* (2–4). A major challenge to eliciting bnAbs is the high sequence variability of the viral glycoprotein Env; accordingly, significant efforts have been made to develop immunogens to focus the antibody response toward a handful of highly conserved regions that correspond to bnAb epitopes (5–8). In contrast to these bnAb epitopes on the native conformation of Env, one highly conserved region of HIV-1 gp41, the N-heptad repeat (NHR), is only transiently exposed during viral entry. Upon binding cellular receptors, Env undergoes substantial conformational changes to form the pre-hairpin intermediate (PHI), in which the NHR and C-heptad repeat (CHR) regions are exposed before the cellular and viral membranes come together for membrane fusion (9–11).

Due to its high sequence conservation (~93%) (12) and critical role in viral entry (9), the NHR is a promising target for blocking HIV-1 infection. Several inhibitors of the NHR have been identified, most notably the FDA-approved fusion inhibitor enfuvirtide (13–15), validating the NHR as a therapeutic target in humans. In addition, cyclic D-peptides that bind the NHR disrupt the fusion process and inhibit HIV-1 infection (16, 17). Of relevance for vaccine applications, several characterized monoclonal antibodies (mAbs) target the NHR (18–20). The first of these mAbs was D5 (18), which was isolated from a human B cell-derived phage display library using two synthetic mimetics of the gp41 PHI: 5-Helix (21) and IZN36 (22). D5 exhibits weak—but broad—neutralizing potency against laboratory-adapted and primary clinical isolates of HIV-1 by binding a conserved hydrophobic pocket of the NHR and preventing the fusion of the viral and cellular membranes (18). X-ray crystallography revealed that complementarity-determining region (CDR) loops in the antibody variable regions of both the heavy and light chains (VH and VL, respectively) contribute to the binding of D5 to the NHR (23). Informed by this insight, Montgomery *et al*. (24) sought to increase the neutralization potency of D5 by randomizing residues in five of the six CDRs (VH CDR 1, 2, 3 and VL CDR 1 and 3). Four D5 IgG variants, each with only one CDR mutated, had slightly increased neutralization potency (24).

Here we hypothesized that combining multiple CDR mutations from these D5 variants would create a more potent mAb against HIV-1. To test this hypothesis, we evaluated a panel of 16 variants: wild-type D5, four previously described CDR variants (24), and 11 recombined CDR variants (Figure 1). The 11 recombined CDR variants are composed of combinations of the CDR sequences found in the four enhanced D5 variants. We determined that D5_AR, the most potent recombined variant, has increased neutralization efficacy in all tier-1 and tier-2 HIV-1 viruses tested compared to D5. As recently reported for D5 (25), and as previously characterized for membrane proximal external region (MPER) mAbs (26, 27), the neutralization potency of D5_AR was enhanced at least 1,000-fold in target TZM-bl cells expressing the high-affinity immunoglobulin receptor FcγRI (TZM-bl/FcγRI cells). Although not expressed on CD4+ T cells, FcγRI is expressed on macrophages and dendritic cells (28), which are thought to be important in the early sexual transmission of HIV-1 at mucosal surfaces (29–36). D5_AR demonstrates cross-clade tier-2 HIV-1 neutralizing activity and extremely potent activity when measured in cells expressing FcγRI, suggesting that the PHI may be a promising vaccine target for eliciting neutralizing antibodies with broad heterologous neutralization potency.

**Figure 1:**
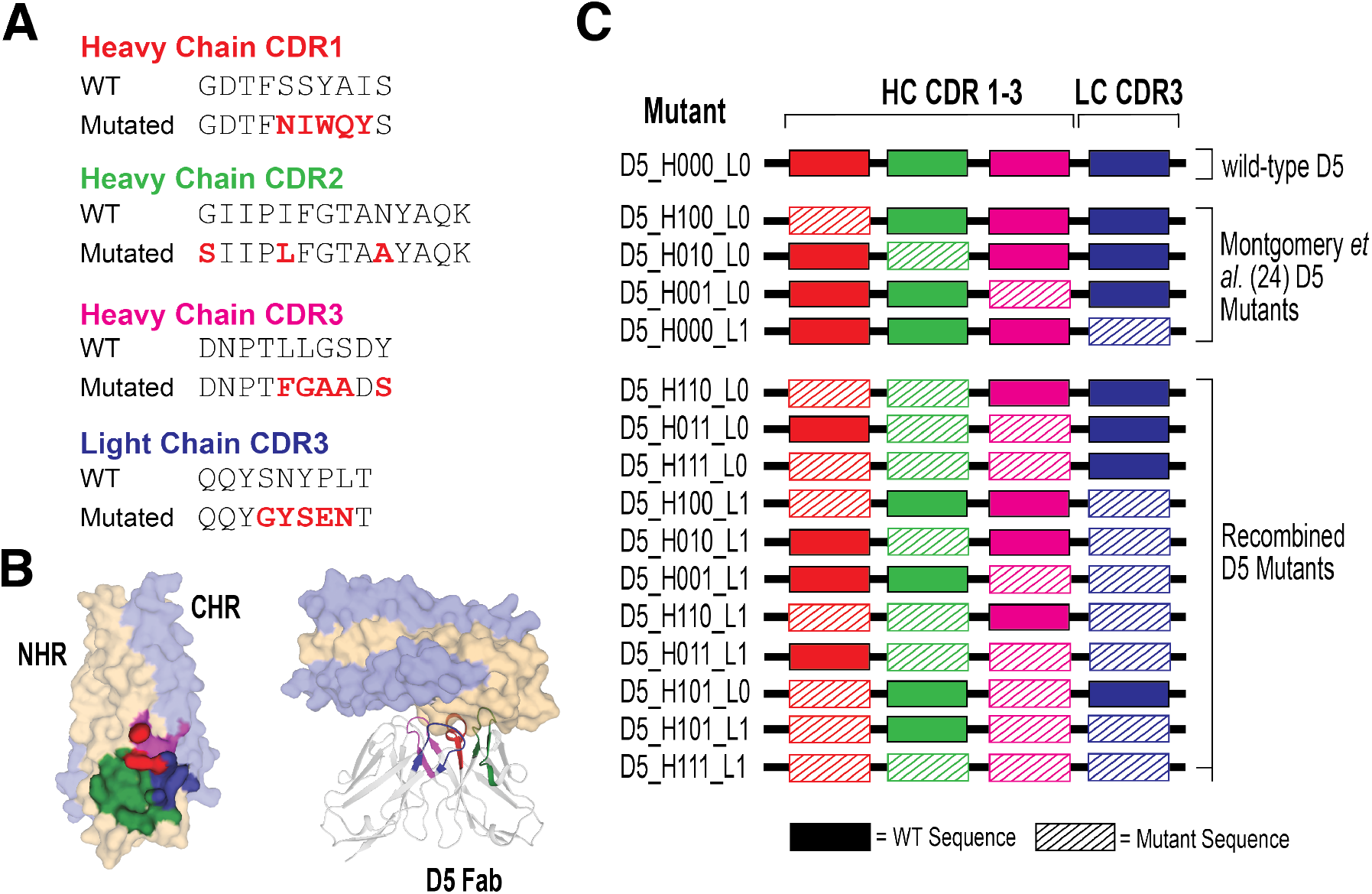
Recombination of known beneficial CDR sequences to engineer novel D5 variants. **(A)** Mutated CDR sequences of four highlighted D5 variants reported by Montgomery *et al*. (24) aligned with the wild-type (WT) sequence. These mutated CDR sequences were recombined to engineer the 11 recombined variants. **(B)** Left: paratope map of the binding sites for D5 Fab with its antigen, 5-Helix. NHR, yellow; CHR, light blue. Areas of contact with the D5 CDR loops are represented as: VH CDR1 (red), VH CDR2 (green), VH CDR3 (magenta), and VL CDR3 (blue). Right: X-ray crystal structure of D5 in complex with 5-Helix (PDB: 2CMR) (23). **(C)** Schematic of D5 variants engineered and tested. The identity of each of the four CDR loops are represented by 0 (WT) or 1 (mutant).

## Results

### D5_AR, a recombined CDR mutant of D5, has enhanced neutralization potency against HIV-1 *in vitro*

To assess the impact on neutralization after the introduction of multiple mutated CDRs, we recombinantly expressed and purified wild-type D5 IgG and 15 D5 IgG variants (Figure 1). The variants were designated D5_HXXX_LX (Figure 1C), with each X replaced by either 0 (representing the wild-type sequence) or 1 (representing the mutated CDR sequence) in the heavy (CDR 1, 2, and 3) or light (CDR3) chain (Figure 1A), reported by Montgomery *et al*. (24). These four CDRs form critical points of contact at the D5 epitope of the NHR (Figure 1B). The full amino acid sequences for the heavy and light chain variable regions of each mutant are provided in Supplemental Figures 1 and 2. Using single-round infectivity assays in TZM-bl cells, each D5 IgG variant was screened for neutralization potency against lentivirus pseudotyped with HIV-1 HXB2 Env (37–41). Neutralization potency was reported as the 50% inhibitory dose (ID_50_). We confirmed that the four single-CDR mutants previously described by Montgomery *et al*. (24) (D5_H100_L0, D5_H010_L0, D5_H001_L0, and D5_H000_L1) displayed enhanced neutralization versus the parent D5 (Supplemental Table 1).

Next, we screened 11 additional D5 variants with lentivirus pseudotyped with HIV-1 HXB2 Env for neutralization potency using a single-round infectivity assay. Several recombinant D5 variants had little effect or even diminished the neutralization potency compared to D5 (Table 1). Nevertheless, we identified six D5 variants that modestly enhanced (>2.0-fold) the neutralization potency of D5 (Table 1). Among these, D5_H011_L0, in which both CDR2 and CDR3 of the heavy chain are mutated, demonstrated the greatest enhancement (4-fold) in ID_50_ (Table 1). We renamed this most potent D5 variant D5_AR for further characterization.

**Table 1:**
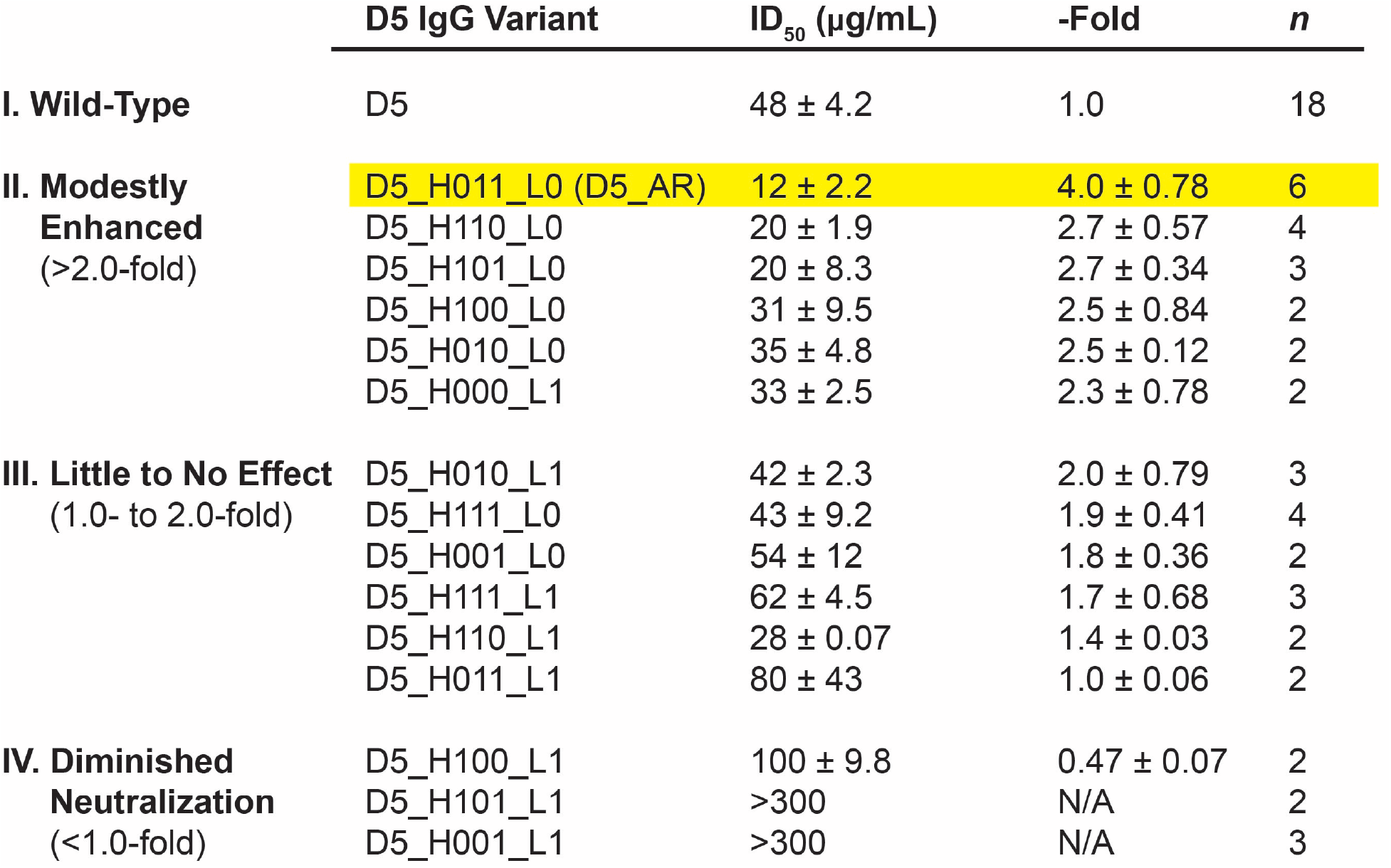
Neutralization profiles of D5 variants. The ID_50_ (half maximal inhibitory dose) of each D5 variant is represented by the geometric mean and standard error of the mean of replicate experiments. For each infection assay, the fold enhancement versus D5 was calculated (ID_50, D5_ / ID_50, D5 variant_); reported fold enhancement is the geometric mean and standard error from replicate experiments. Fold enhancement >1 corresponds to enhanced neutralization potency (reduced ID_50_).

Our initial neutralization screen utilized antibody purified via Protein A affinity chromatography (Materials and Methods). During subsequent antibody purification, we removed aggregates through an additional size exclusion chromatography (SEC) step; comparison of neutralization potencies revealed that the presence of aggregates reduced the neutralization potencies of D5 and D5_AR (Supplementary Information, Supplemental Figure 3). Experiments in Table 1 and Supplemental Table 1 were the only experiments reported here that used non-SEC-purified antibody preparations, which accounts for the difference in ID_50_ values for HXB2 reported in Table 1 and Figure 3.

### D5 and D5_AR neutralize with similar potency as an scFv, Fab, and IgG

In the first description of D5, the IgG (~150 kD) and scFv (~25 kD) constructs neutralized similarly to one another (18). However, more recent studies reported that D5 scFv is more potent than D5 Fab (~50 kD), and that both were more potent than D5 IgG (20, 23, 24, 42); in addition, there are reports that increasing the size of NHR inhibitors reduces neutralization potency (20, 24, 42–44). These size-dependent findings would imply steric hindrance in accessing the PHI. Given our finding that SEC purification could impact the neutralization potency of D5 and D5_AR IgG (Supplemental Figure 3A), we decided to reinvestigate the question of size-dependent neutralization for D5 using antibody preparations free of aggregates. Notably, in agreement with the initial report (18) and in contrast to the previous reports (20, 23, 24, 42), we found that D5 as an IgG, Fab, and scFv (all SEC-purified) did not exhibit a size-dependent pattern of neutralization (Figure 2A). Additionally, we detected comparable neutralization potency for D5_AR as an scFv, Fab, and IgG (Figure 2B). These results demonstrate that neither D5 nor D5_AR are impacted by steric hindrance and suggest the presence of protein aggregates could explain previous reports of size-dependent neutralization for D5 scFv, Fab, and IgG.

**Figure 2:**
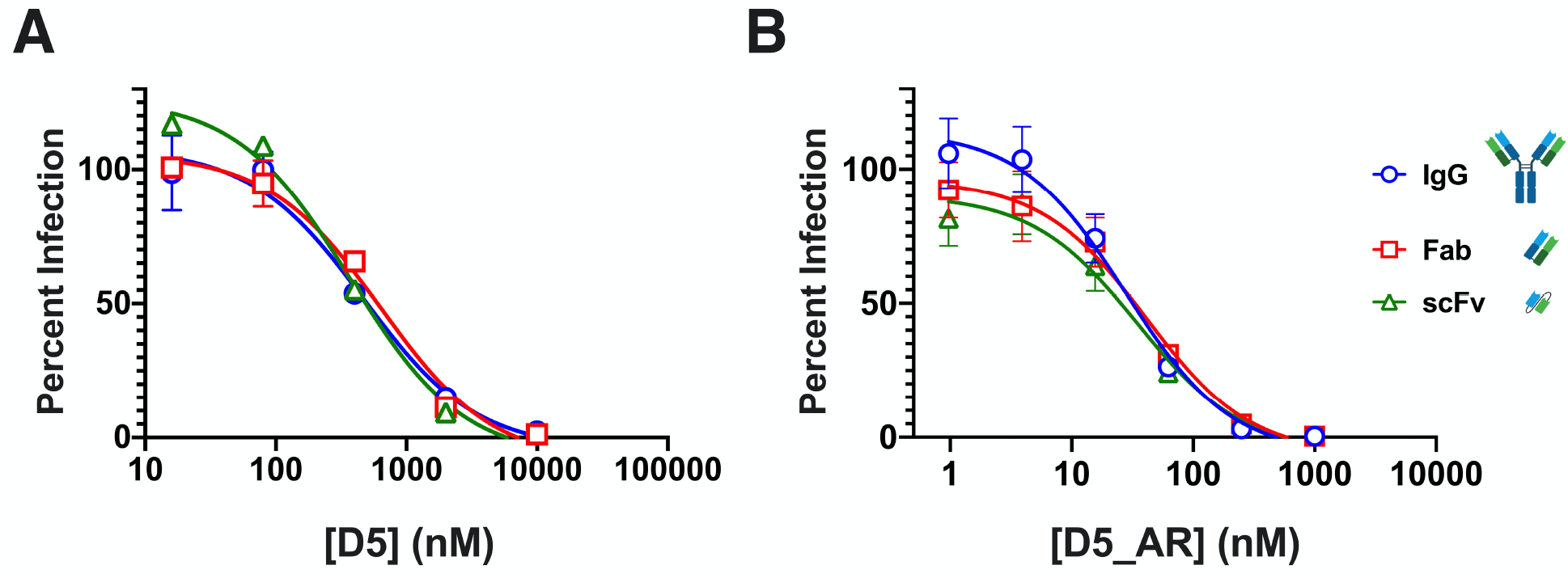
D5 and D5_AR neutralize HIV-1 *in vitro* in a size-independent manner. scFv, Fab, and IgG constructs of **(A)** D5 and **(B)** D5_AR are similarly effective *in vitro* at neutralizing lentivirus pseudotyped with HIV-1 HXB2. Data points and error bars are the mean percent infection and standard error of the mean, respectively (n=2). Antibody construct images used in this figure were generated with BioRender.

### D5_AR exhibits more potent cross-clade tier-2 neutralization of HIV-1 viruses than D5

We next investigated the potency of D5_AR IgG in neutralizing a diverse panel of 19 tier-1 and tier-2 pseudotyped viruses across eight viral clades (A, AC, B, C, G, CRF01, CRF02, and CRF07) (40, 45–60). Ten of these strains originated from a 12-virus panel designed to capture the sequence diversity of the HIV-1 epidemic globally (40). Tier 1A and tier 1B contain viruses that are most sensitive to neutralization by antibodies, whereas tier-2 viruses have modest sensitivity to neutralizing antibodies (52). D5_AR IgG neutralized virus more effectively than D5 IgG across all tiers and clades (Figure 3A). Notably, D5_AR neutralizes more viruses with an ID_50_ of <50 μg/mL (63%) than D5 (11%) (Figure 3B).

**Figure 3:**
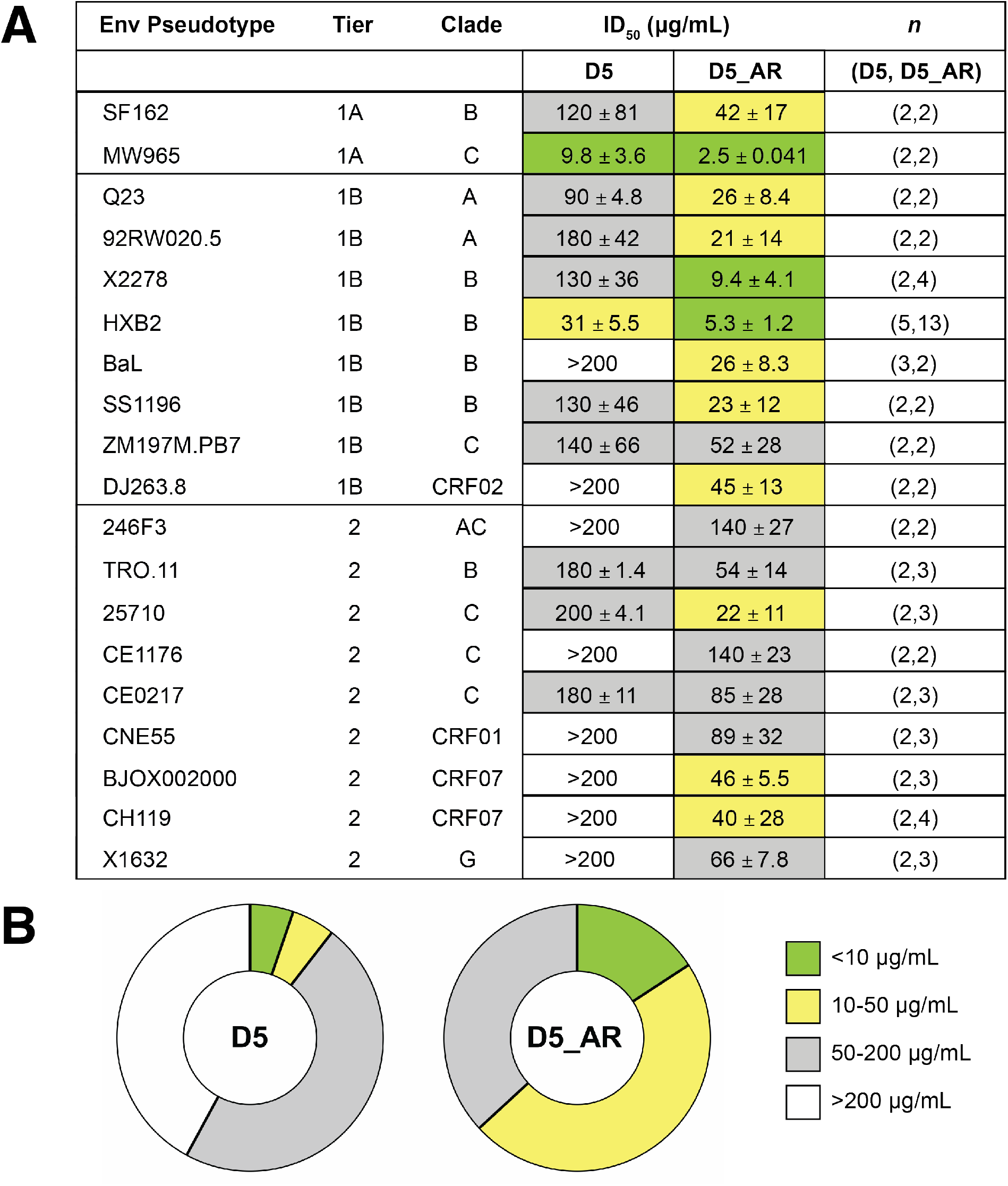
D5_AR IgG has higher neutralization potency than D5 across tiers and clades of HIV-1. **(A)** D5_AR IgG has an enhanced neutralization profile versus D5 IgG, as indicated by ID_50_ values (geometric mean ± standard error of the mean) against a panel of tier-1 and tier-2 HIV-1 viruses from multiple clades. **(B)** D5_AR IgG neutralizes a greater percentage of HIV-1 viruses than D5 IgG within the <10 μg/mL and 10-50 μg/mL ID_50_ ranges.

### D5_AR neutralization potency is enhanced >1,000-fold in FcγRI-expressing cells

Recently, we reported the neutralization potency of D5 IgG to be greatly increased in TZM-bl cells expressing the cell-surface receptor FcγRI (TZM-bl/FcγRI cells) (25). FcγRI is the only known high-affinity IgG receptor in humans, capable of binding monomeric IgG (61). To determine whether D5_AR IgG is similarly potentiated, we tested its neutralization of an additional panel of tier-2 HIV-1 viruses and SHIV challenge viruses in TZM-bl/FcγRI cells (Figure 4). Indeed, neutralization by D5_AR IgG was potentiated ~1,000-fold in TZM-bl/FcγRI cells versus TZM-bl cells (Figure 4A and 4B). In the presence of FcγRI, D5_AR had potent neutralization activity against a panel of tier-2 HIV-1 viruses, with ID_50_ values <0.1 μg/mL (Figure 4C and 4D). Consistent with an Fc-dependent mechanism, the Fab form of D5_AR did not exhibit potentiation (Figure 5A). This observed potentiation was specific to FcγRI: enhanced neutralization was minimal or not observed in cell lines expressing other Fc receptors (FcγRIIa, FcγRIIb, and FcγRIIIa; Figure 5B). It is noteworthy that the ID_50_ values of D5_AR IgG in TZM-bl cells were approximately linearly related to the ID_50_ values in TZM-bl/FcγRI cells (Figure 6).

**Figure 4:**
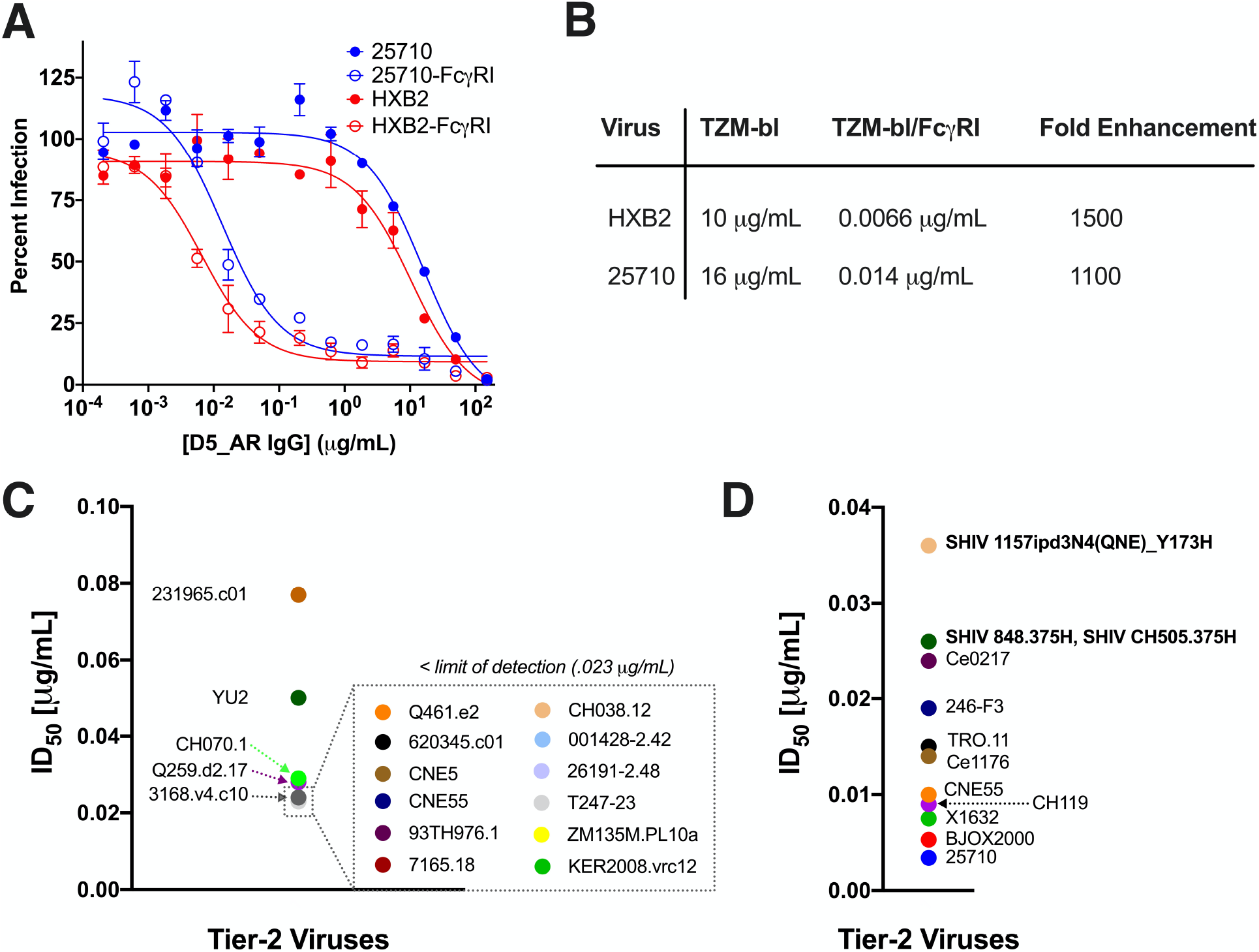
The neutralization potencies of D5_AR IgG against tier-2 HIV-1 viruses are substantially higher in TZM-bl cells expressing FcγRI. **(A)** Neutralization curves demonstrating the enhanced potency of D5_AR IgG against lentivirus pseudotyped with HIV-1 HXB2 (tier 1B) and 25710 (tier 2) in TZM-bl cells expressing FcγRI versus TZM-bl cells without FcγRI. Data points and error bars are mean and standard error of the mean (n=2), respectively. **(B)** ID_50_ values and fold enhancement of neutralization potency of D5_AR IgG against lentivirus pseudotyped with HIV-1 HXB2 (tier 1B) and 25710 (tier 2) in TZM-bl cells versus TZM-bl/FcγRI cells. D5_AR is potentiated approximately 1,000-fold in TZM-bl/FcγRI cells. **(C)** ID_50_ values for D5_AR against a panel of tier-2 HIV-1 viruses in TZM-bl/FcγRI cells. Many of the viruses had ID_50_ values below the limit of detection (0.023 μg/mL). **(D)** ID_50_ values for D5_AR against another panel of tier-2 and SHIV challenge viruses in TZM-bl/FcγRI cells but with a lower limit of detection.

**Figure 5:**
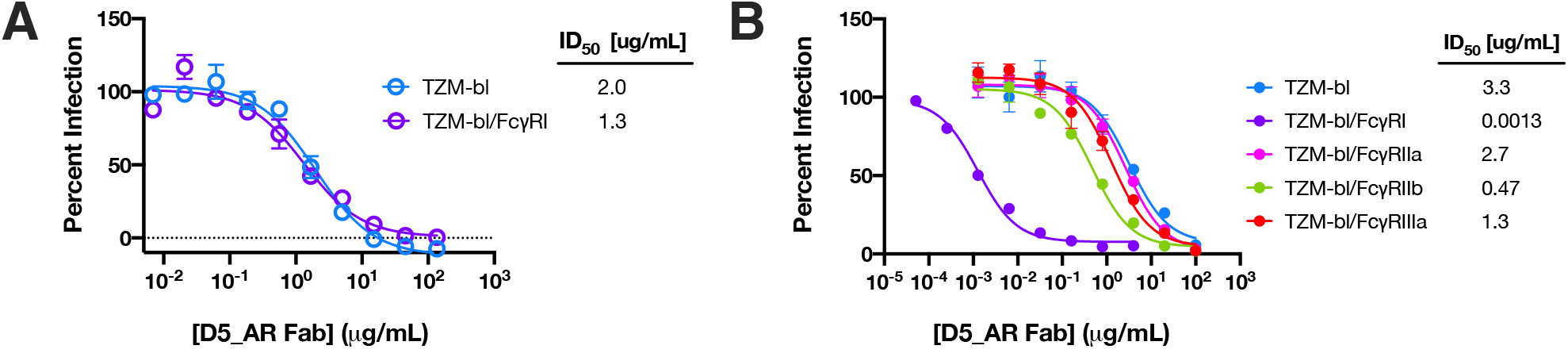
D5_AR is not potentiated as a Fab and shows minimal or no potentiation in the presence of other Fc receptors. **(A)** Neutralization curves for D5_AR Fab against lentivirus pseudotyped with HIV-1 HXB2 show no neutralization enhancement between TZM-bl cells and TZM-bl/FcγRI cells. Data points and error bars are the mean and standard error of the mean (n=2), respectively. **(B)** The degree of observed neutralization enhancement is FcγRI-specific, as demonstrated by the neutralization potency of D5 AR IgG against lentivirus pseudotyped with HIV-1 HXB2 in TZM-bl cells expressing various Fc receptors. Data points and error bars are the mean and standard error of the mean (n=3), respectively.

**Figure 6:**
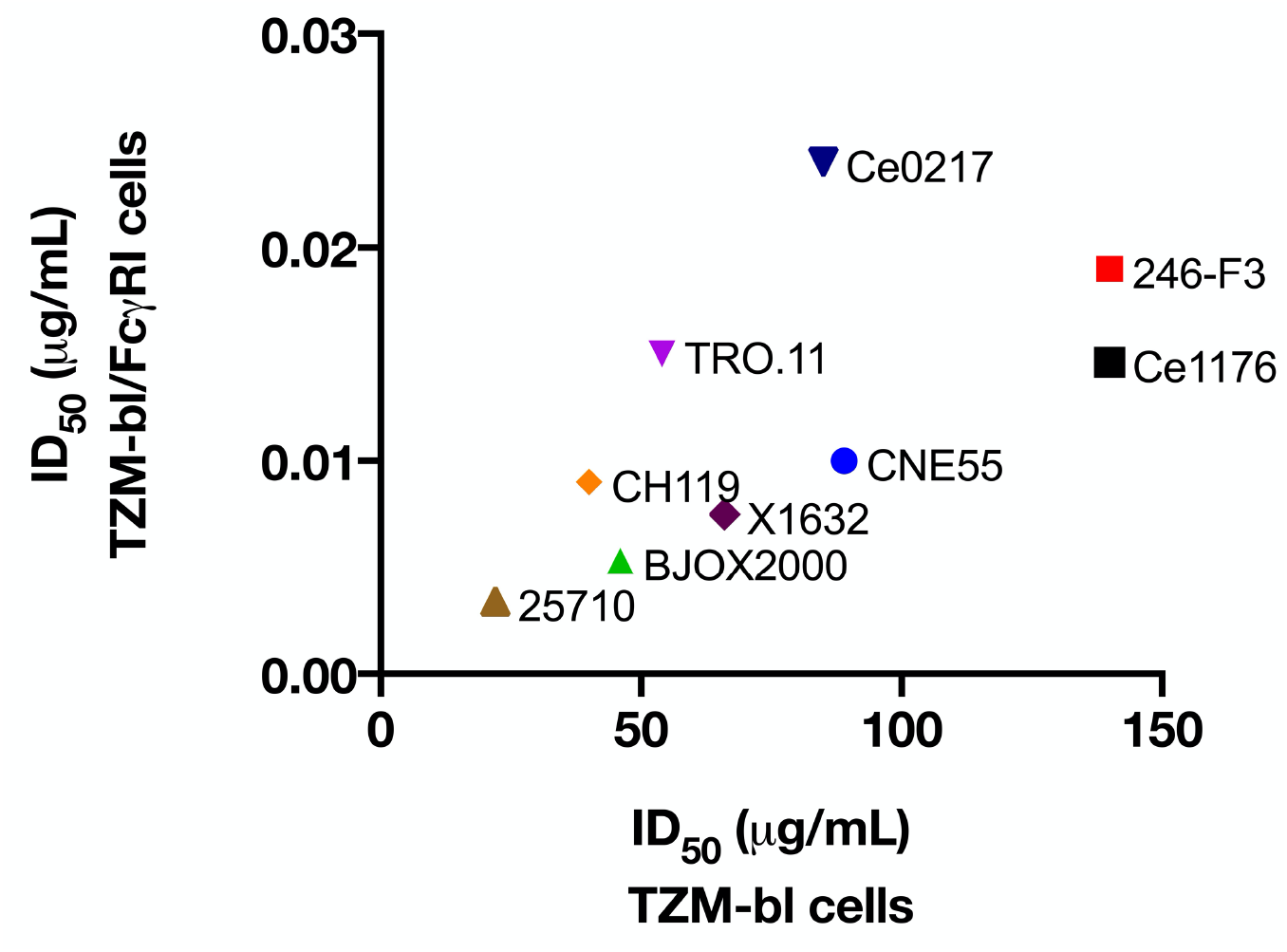
Comparison of HIV-1 neutralizing activity of D5 AR IgG in TZM-bl versus TZM-bl/FcγRI cells. The ID_50_ values of D5 AR IgG in TZM-bl and TZM-bl/FcγRI cells are approximately linearly related in a panel of tier-2 HIV-1 viruses from multiple clades. ID_50_ values represented in this figure were reported previously in Figure 3 and 4.

## Discussion

The PHI of HIV-1 is a validated drug target in humans (13–15), and antibodies that bind the NHR of gp41 that is exposed in the PHI can inhibit HIV-1 infection *in vitro* (18–20). The first of these NHR-binding antibodies, D5, has weak neutralization potency against tier-1 HIV-1 strains (18). Here we engineered and characterized a more potent D5 derivative, D5_AR, which exhibited enhanced neutralization of tier-1 strains and weak cross-clade neutralization of tier-2 HIV-1 viruses (Figure 3). Using ten global HIV-1 reference strains, we determined that at modest concentrations (1-50 μg/mL), D5_AR neutralizes the majority of tier-1 and tier-2 HIV-1 strains tested from a variety of clades (Figure 3). Taken together with the high sequence conservation of the NHR (12), these results support future vaccine design efforts to elicit neutralizing antibodies against the PHI.

Several previous reports had suggested that access to the D5 epitope was impacted by steric hindrance, as smaller antibody constructs were more potent than full-length IgG (20, 23, 24, 42). We reinvestigated this issue using antibody preparations that were free of observed protein aggregates (Supplementary Information; Supplemental Figure 3). In contrast to earlier reports (20, 23, 24, 42), here we confirm (18) that D5 and D5_AR are similarly potent when tested in neutralization assays in scFv, Fab, and IgG formats (Figure 2). We hypothesize that the presence of higher-order protein aggregates (that can be removed by SEC) may explain the previous reports of size-dependent neutralization by D5. Given these findings for D5 and D5_AR, we conclude that steric hindrance is not an obstacle for at least some anti-PHI antibodies.

Previous work on antibodies targeting another epitope of gp41, the MPER, found that neutralization activity was potentiated as much as 5,000-fold in cells expressing FcγRI, an integral membrane protein that interacts with the Fc portion of γ immunoglobulins (26–28). Since the MPER is not fully exposed until after Env engages with cellular receptors (62, 63), these results suggest that by binding the Fc region of MPER mAbs, FcγRI provides a local concentration advantage at the cell surface that enhances neutralization (26, 27). Because the PHI, like the MPER, is fully exposed only during viral membrane fusion, we previously investigated the effect of FcγRI on D5 and found neutralization by this anti-PHI mAb is also enhanced ~5,000-fold (25). Like D5, D5_AR IgG displayed ~1,000-fold enhancement in neutralization potency in TZM-bl/FcγRI cells (Figure 4A and 4B). Notably, this enhancement makes D5_AR IgG an extremely potent neutralizing antibody of tier-2 HIV-1 viruses in the TZM-bl/FcγRI cell line, with ID_50_ values below 0.1 μg/mL (Figure 4C and 4D).

TZM-bl/FcγRI cells enable sensitive detection of neutralization activity from anti-NHR antibodies and could be used, in conjunction with TZM-bl cells, to monitor progress toward eliciting neutralizing antisera. We hypothesize that neutralizing activity detected by TZM-bl/FcγRI cells could be used as an indicator of low-affinity antibody precursors in serum that have the potential to mature to high-affinity neutralizing activity independent of FcγRI. This hypothesis is supported by our findings that the neutralizing activity of D5_AR in FcγRI-expressing cells were approximately linearly related to the neutralizing activity of D5_AR in cells without FcγRI (Figure 6).

Although not normally expressed on CD4+ T cells, FcγRI receptors are expressed on some cells at mucosal surfaces where sexual HIV-1 transmission occurs, such as macrophages and dendritic cells (28). Mucosal macrophages and dendritic cells can be productively infected by HIV-1 (29–32) and can then mediate viral transmission to CD4+ T cells (33–36). Importantly, studies of intravaginal inoculation of simian immunodeficiency virus (SIV) of non-human primates demonstrated that intraepithelial and submucosal dendritic cells are infected in the earliest stages (18-48 hours) of SIV infection (64–67). More recent work has shown that at 48 hours post-inoculation, 25% of infected cells are dendritic cells and macrophages, with the remainder comprising CD4+ T cells, primarily of the Th17 type (68, 69). Thus, it is plausible that inhibiting HIV-1 infection of these FcγRI-expressing cells at the mucosal surfaces could decrease the likelihood of sexual HIV-1 transmission.

Indeed, in a vaginal challenge with SHIV in rhesus macaques, an MPER mAb (2F5) afforded dose-dependent protection when administered as an IgG, but not when administered in its Fab form (70), suggesting an Fc-dependent mechanism of protection *in vivo*. Previous studies have also demonstrated that MPER mAbs are much more protective against SHIV challenge than when measured *in vitro*, compared to other bnAbs (71–73). Considered alongside these previous findings, the extremely potent activity of D5_AR observed here against HIV-1 infection of FcγRI-expressing cells (Figure 4) motivates future efforts to investigate the ability of passively transferred anti-PHI antibodies to protect against sexual HIV-1 transmission *in vivo*.

Taken together, the findings presented here provide evidence that the NHR of the PHI is a promising target for future HIV-1 vaccine development, and pave the way for future studies of the *in vivo* significance of FcγRI-mediated potentiation of anti-PHI antibodies.

## Materials and Methods

### Mammalian expression of D5 constructs

The variable heavy (VH) chain regions of the D5 variants were ordered as gene fragments from Twist Biosciences (Supplemental Figure 1A). Gene fragments were resuspended to 10 ng/μL in H_2_O and PCR amplified using the following two primers: (HC_forward) 5’-ACCGGTGTACATTCCCAGGTTCAAC-3’ and (HC_reverse) 5’-GCCCTTGGTCGACGCGCTTGATACG-3’. The mutated variable light (VL) chain region, with only the third CDR mutated according to Montgomery *et al*. (24), was ordered as a G-block gene fragment from Integrated DNA Technologies and PCR amplified using the following two primers: (LC_forward) 5’-ACCGGTGTACATTCAGATATTCAAATGAC-3’ and (LC_reverse) 5’ - TGCAGCCACCGTACGTTTG-3’. Purified VH and VL fragments were cloned into linearized pCMVR with either the human IgG heavy or kappa light constant regions, respectively (74, 75). The primers for linearizing the pCMVR IgG heavy chain plasmid were: (HC_lin_forward) 5’-GCGTCGACCAAGGGCCCATCGGTCTTC-3’ and (HC_lin_reverse) 5’-GGAATGTACACCGGTTGCAGTTGCTACTAGAAAAAG-3’. The primers for linearizing the pCMVR IgG kappa light chain plasmid were: (LC_lin_forward) 5’-CGTACGGTGGCTGCACCATCTGTCTTCATCTTC-3’ and (LC_lin_reverse) 5’-TGAATGTACACCGGTTGCAGTTGCTACTAGAAAAAGGATGATA-3’. The D5 VH and VL segments were cloned into the linearized pCMVR backbones with 5X In-Fusion HD Enzyme Premix (Takara Bio). Plasmids were transformed into Stellar Competent Cells (Takara Bio) and transformed cells were grown at 37 °C. Colonies were sequence confirmed and then maxi-prepped (NucleoBond^®^ Xtra Maxi, Macherey-Nagel). Plasmids were sterile filtered using a 0.22-μm syringe filter and stored at −20 °C.

D5 IgG variants used for neutralization assays were expressed in Expi293F™ cells (Thermo Fisher Scientific) using FectoPRO^®^ (Polyplus). VH and VL plasmids were co-transfected at a 1:2 ratio, respectively; cells were transfected at 3×10^6^ cells/mL. Cell cultures were incubated at 37 °C and 8% CO_2_ with shaking at 120 rpm. Cells were harvested 3 days post transfection by spinning at 300 x *g* for 5 min and then filtered through a 0.22-μm filter. IgG supernatants were diluted 1:1 with 1X phosphate-buffered saline (PBS) and batch-bound to Pierce™ Protein A agarose (Thermo Fisher Scientific) overnight at 4 °C. The supernatant/resin slurry was added to a column and the resin was washed with 1X PBS and eluted with 100 mM glycine [pH 2.8] into 1/10 volume of 1 M Tris [pH 8.0].

D5 and D5_AR Fab used for neutralization assays were also produced in Expi293F™ cells. D5 and D5_AR Fab VH regions were cloned into a pCMVR heavy chain linearized backbone with a portion (CH2 and CH3 domains) of the constant region removed. Fab VH and VL plasmids were co-transfected and harvested with the protocol for IgG described above. Fab supernatants were diluted 1:1 with 50 mM sodium acetate [pH 5.0], batch-bound to Pierce™ Protein G Agarose (Thermo Fisher Scientific) overnight at 4 °C, washed with 50 mM sodium acetate [pH 5.0], and eluted with 100 mM glycine [pH 2.8] into 1/10 volume of 1 M Tris [pH 8.0].

D5 and D5_AR scFv constructs used for neutralization assays were expressed in Expi293F™ cells. The VH and VL regions were linked via a human muscle aldolase sequence (76), tagged with a His6-tag, and cloned into a linearized pCMVR vector (Supplemental Figure 2). The scFv plasmid was transfected and harvested with the same protocol as IgG and Fab. ScFv supernatants were diluted 1:1 with 10 mM imidazole in 1X PBS, batch-bound to Ni-NTA agarose (Thermo Fisher Scientific) overnight at 4 °C, washed with 10 mM imidazole in 1X PBS, and eluted with 250 mM imidazole in 1X PBS.

### Purification and storage of D5 constructs

For the initial screening in neutralization assays (Table 1 and Supplemental Table 1), there was no purification following elution from Protein A affinity purification. Elutions were buffer exchanged and spin concentrated using 1X PBS and Amicon^®^ Ultra-15 10 kD 15 mL spin concentrators (Millipore).

For all other neutralization assays, elutions were further purified after affinity purification on an AKTA™ using a GE Superdex 200 Increase 10/300 GL column (GE HealthCare) in 1X PBS. After size exclusion chromatography, samples were spin concentrated using Amicon^®^ Ultra-15 10 kD 15 mL spin concentrators.

Fab and scFv constructs were eluted from affinity purification and then purified further via size exclusion chromatography using the Superdex 200 Increase 10/300 GL column (GE HealthCare) and 1X PBS. Samples were spin concentrated as described above.

For all samples, regardless of the purification procedure, concentrated elution samples were syringe filtered using a 0.22-μm filter and stored at 4 °C prior to use.

### Transfection to produce HIV-1 pseudotyped lentiviruses

HEK293T cells were transiently co-transfected with a backbone plasmid as well as a HIV-1 Env plasmid for HIV-1 pseudotyped lentivirus production using the calcium phosphate transfection protocol previously described (77–79). HEK293T cells were passaged in T75 flasks and incubated at 37 °C at 5% CO_2_. The growth medium used for passaging and transfections was Corning^®^ DMEM (Dulbecco’s Modified Eagle Medium with 4.5 g/L glucose, l-glutamine, and sodium pyruvate) with 10% fetal bovine serum, 1% penicillin streptomycin (Corning), and 1% l-glutamine (Corning).

The backbone plasmid psg3ΔEnv was obtained through the NIH AIDS Reagent Program, Division of AIDS, NIAID, NIH from Drs. John C. Kappes and Xiaoyun Wu: HIV-1 SG3 ΔEnv Non-infectious Molecular Clone (Cat#11051) (80, 81). The psg3ΔEnv plasmid was propagated in MAX Efficiency^®^ Stbl2™ cells grown at 30 °C with shaking and Env plasmids were propagated in Stellar Competent Cells grown at 37 °C with shaking. DNA was isolated using a maxi-prep kit (NucleoBond^®^ Xtra Maxi, Macherey-Nagel) and sequence confirmed.

In brief, 6×10^6^ HEK293T cells were plated in 10-cm petri dishes in a total volume of 10 mL of DMEM and incubated overnight at 37 °C and 5% CO_2_ without shaking. Once the cells reached 50-80% confluency, they were transfected as follows. In a Falcon tube, 20 μg of psg3ΔEnv was mixed with 10 μg of Env plasmid and water for a final volume of 500 μL. Five hundred microliters of 2X HEPES-buffered saline [pH 7] (Alfa Aesar) were added dropwise to the mixture and 100 μL 2.5 M CaCl2 were added subsequently. The mixture was incubated at room temperature for 20 min and then added dropwise onto the cells. Next, 12-18 h after transfection, the medium was aspirated from the dish and replaced with 10 mL of fresh DMEM with additives. Virus-containing medium was harvested 48 h after medium swap and centrifuged at 300 x *g* for 5 min; the supernatant was sterile-filtered with a 0.45-μm polyvinylidene difluoride filter and stored in 1-mL aliquots at −80 °C.

### Neutralization Assay

The neutralization assay was adapted from the TZM-bl assay protocol using HIV-1 Env-pseudotyped viruses as described previously (38, 41). Briefly, TZM-bl cells, derived from the JC53-bl parental cell line, were used as reporter cells in this assay and were obtained through the NIH AIDS Reagent Program (Cat#8129) from Dr. John C. Kappes, and Dr. Xiaoyun Wu (80, 82–85). TZM-bl cells are adherent HeLa cells that stably express CD4 and CCR5 and constitutively express CXCR4; they have integrated β-galactosidase and firefly luciferase reporter genes under the control of the HIV-1 LTR promoter. TZM-bl cells transduced to stably express FcγRI (26, 27), were also used in these neutralization assays. TZM-bl cells were passaged in T25 flasks and incubated at 37 °C at 5% CO_2_ without shaking. The growth medium used for passaging and neutralization assays was Corning^®^ DMEM with 10% fetal bovine serum, 1% penicillin streptomycin (Corning), and 1% l-glutamine (Corning).

In brief, 5×10^3^ TZM-bl cells were plated in the internal 60 wells of white-walled, clear-bottom, 96-well plates and incubated overnight at 37 °C, 5% CO_2_ without shaking. All outside wells were filled with 200 μL PBS to minimize evaporation. On the next day, the medium was aspirated without disturbing the cells and replaced with a final mixture composed of ¼ volume DMEM, ½ volume HIV-1 pseudotyped lentivirus, ½ volume D5 antibody at varying concentrations, and DEAE dextran (10 μg/mL). Forty-eight hours after infection, all medium was aspirated off the wells, cells were lysed and either luciferase activity was determined using BriteLite Plus Reagent (Perkin Elmer) or β-galactosidase activity was determined using Tropix Gal-Screen™ (Applied Biosystems) and Buffer A (Applied Biosystems). β-galactosidase readout was used for neutralizations show in Table 1, Figures 2 and 3. Luciferase readout was used for neutralizations shown in Figures 4 and 5.

Relative luminescent unit (RLU) values were quantified using a Synergy™ HTX Multi-Mode Reader (BioTek^®^), normalized against cells-only reference wells, and averaged for technical replicates on the plate. Percent infectivity and propagated error values (Statistics and Data Analysis) were entered into GraphPad Prism 8. Neutralization titers are reported as the antibody concentration at which RLU were reduced by 50% compared to RLU in virus control wells after subtraction of background RLU in cell control wells. ID_50_ was calculated using the [inhibitor] vs. response (three parameters) dose-response curve fit in GraphPad Prism 8. This assay was conducted in compliance with good clinical laboratory procedures (86), including participation in a formal TZM-bl assay proficiency program for GCLP-compliant laboratories (37).

### Statistics and Data Analysis

Percent infectivity for the neutralization assays was calculated as follows: 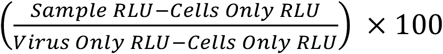. Propagated error for the percent infectivity was calculated using the following formula: 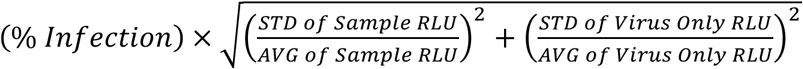. The ID_50_ values in Table 1 and Figure 3 represent the geometric mean of the biological replicates for the tested antibodies with the standard error of the mean reported. Fold difference in ID_50_ was calculated for each experiment by dividing the D5 ID_50_ by the D5 variant ID_50_. Because fold difference was calculated for each experiment, the reported fold differences in Table 1 and Figure 3 are the geometric mean and the standard error of the mean from all replicates.

## Author Contributions

AAR, MVFI, CLB, and PSK conceived the experiments. AAR performed protein purification, antibody characterization, and neutralization assays. MVFI and BNB performed protein purification and neutralization assays. Additional neutralization assays were also conducted under the supervision of CCL and DCM. All authors contributed to revising the manuscript.

## Acknowledgments

We thank members of the Kim Lab for helpful discussions and Drs. A. E. Powell, S. Tang, and D. Xu for critical reading of this manuscript. This research was supported by the National Institute of General Medical Sciences of the National Institutes of Health under award numbers T32GM007276 (BNB) and 5T32GM007365 (MVFI), the NSF GRFP (BNB), NIH/NIAID contract HHSN272201800004C (CCL), the Bill and Melinda Gates Foundation award OPP1113682 (PSK), the Virginia and D. K. Ludwig Fund for Cancer Research (PSK), and the Chan Zuckerberg Biohub (PSK).

## Supplementary Information

### Purification method impacts D5’s neutralization potency

After completing the initial neutralization screens of the D5 variants, a more in-depth neutralization profile was determined for D5_AR compared to D5. This profiling involved testing the constructs’ neutralization potency against various viruses and in various antibody formats. Notably, we found that the manner of purification and concentration of antibodies impacts the observed neutralization activity of the antibody. In our initial neutralization screens, we did not include a size exclusion chromatography (SEC) purification step after Protein A purification: we immediately spin-concentrated the elution and filtered the concentrated antibody for use in neutralization assays (Materials and Methods). For the in-depth neutralization analysis, we further purified antibodies via SEC after Protein A purification. After SEC purification, we concentrated the protein using spin concentrators. These ID_50_ values (Figure 3) differed from the values that we previously measured (Table 1) when the antibodies were not SEC-purified (SEC-purified preparations had a 1.5-fold decrease in ID_50_ for D5 and a 2.3-fold decrease in ID_50_ for D5_AR) (Supplemental Figure 3). A side-by-side comparison revealed that there was indeed a difference in neutralization for D5-AR (Supplemental Figure 3A), although neutralization was still enhanced compared to D5. UV traces from SEC indicated that aggregates may explain this difference in reported neutralization (Supplemental Figure 3B), which likely explains the discrepancy in reported ID_50_ values for HXB2 in Table 1 and Figure 3 of the main text.

### Supplemental Figures and Tables

**Supplemental Figure 1:**
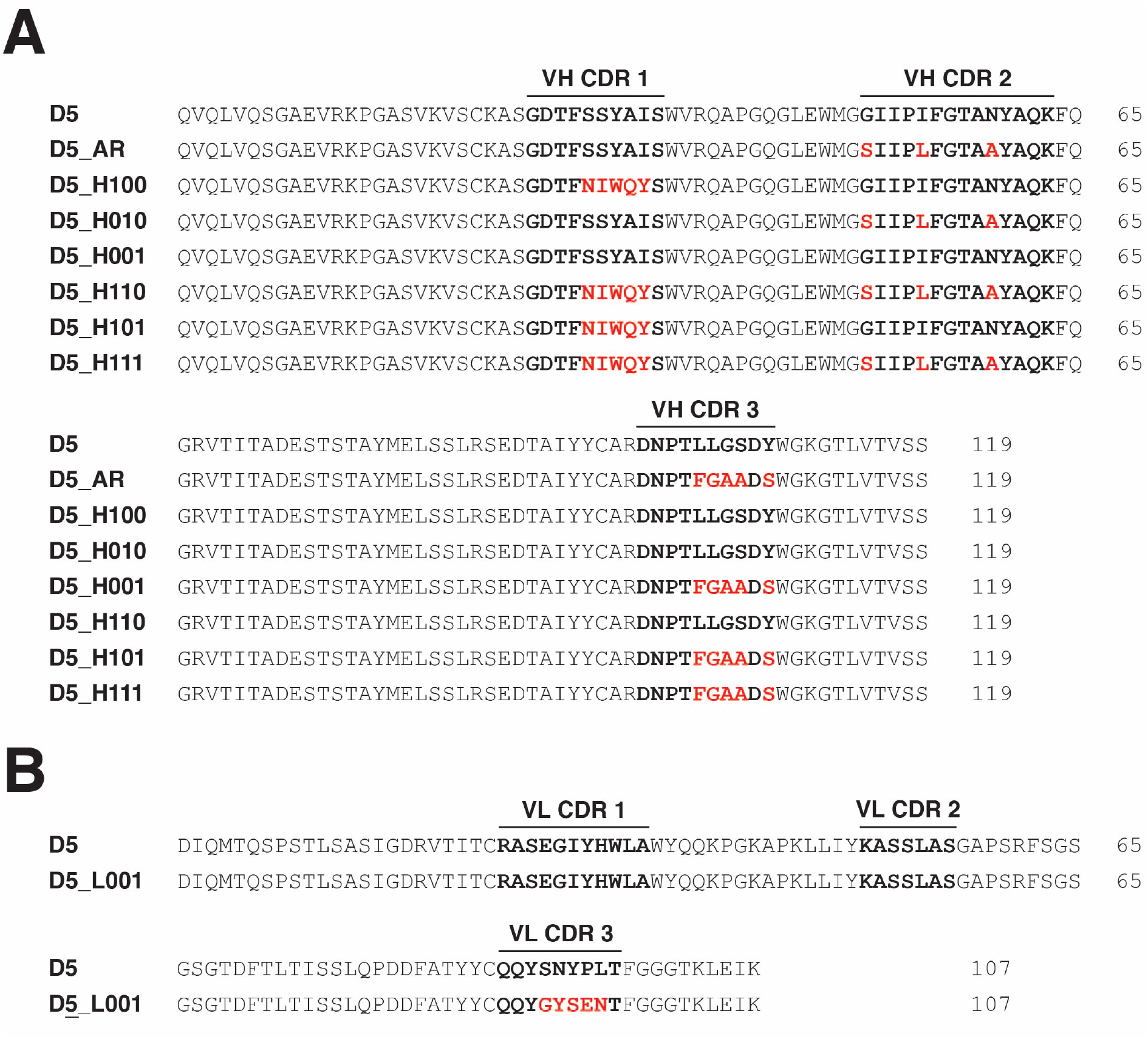
Sequences of D5 variant variable heavy and variable light chain regions. **(A)** Sequence alignment of variable heavy (VH) chain sequences of D5 variants (IGHV1-69 germline). **(B)** Sequence alignment of variable light (VL) chain sequence of D5 variants (IGKV1-5 germline). Bold font denotes complementarity-determining regions and red letters highlight amino acid changes from wild-type D5.

**Supplemental Figure 2:**
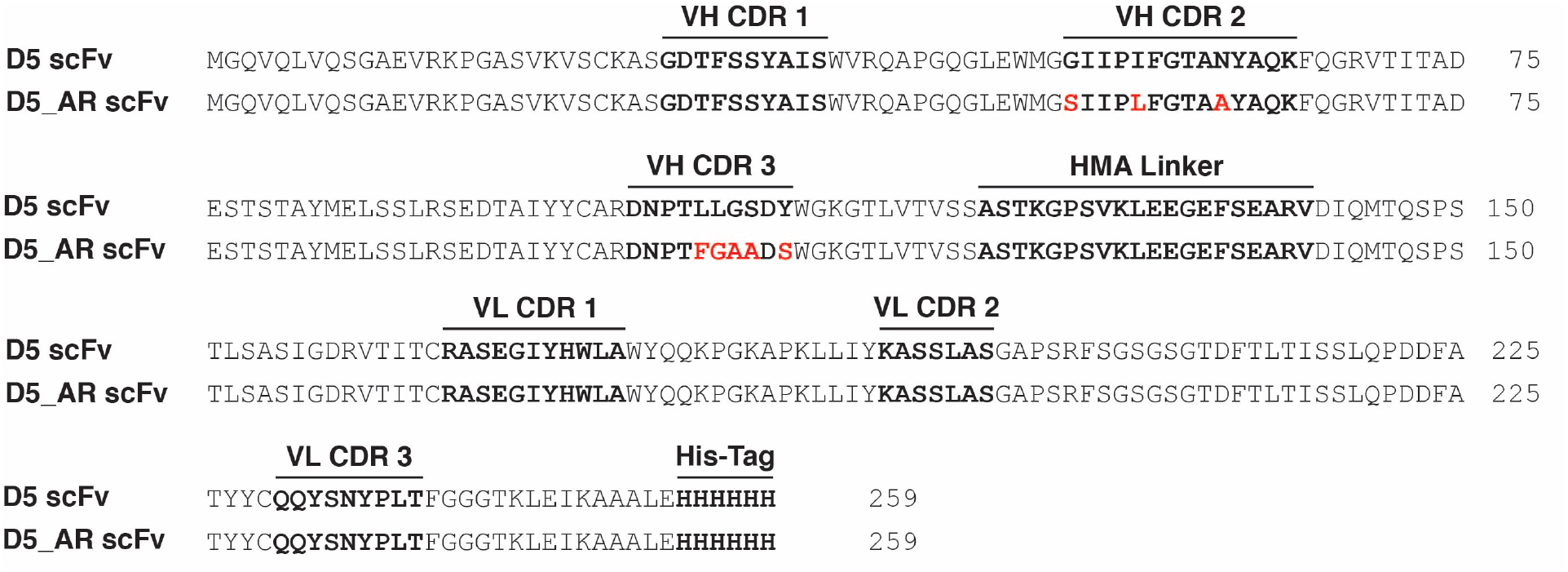
Sequences of D5 and D5_AR scFv inserts with human muscle aldolase linker and His6-Tag. Bold font denotes complementarity-determining regions, the linker, and the tag; red letters highlight amino acid changes from wild-type D5.

**Supplemental Figure 3:**
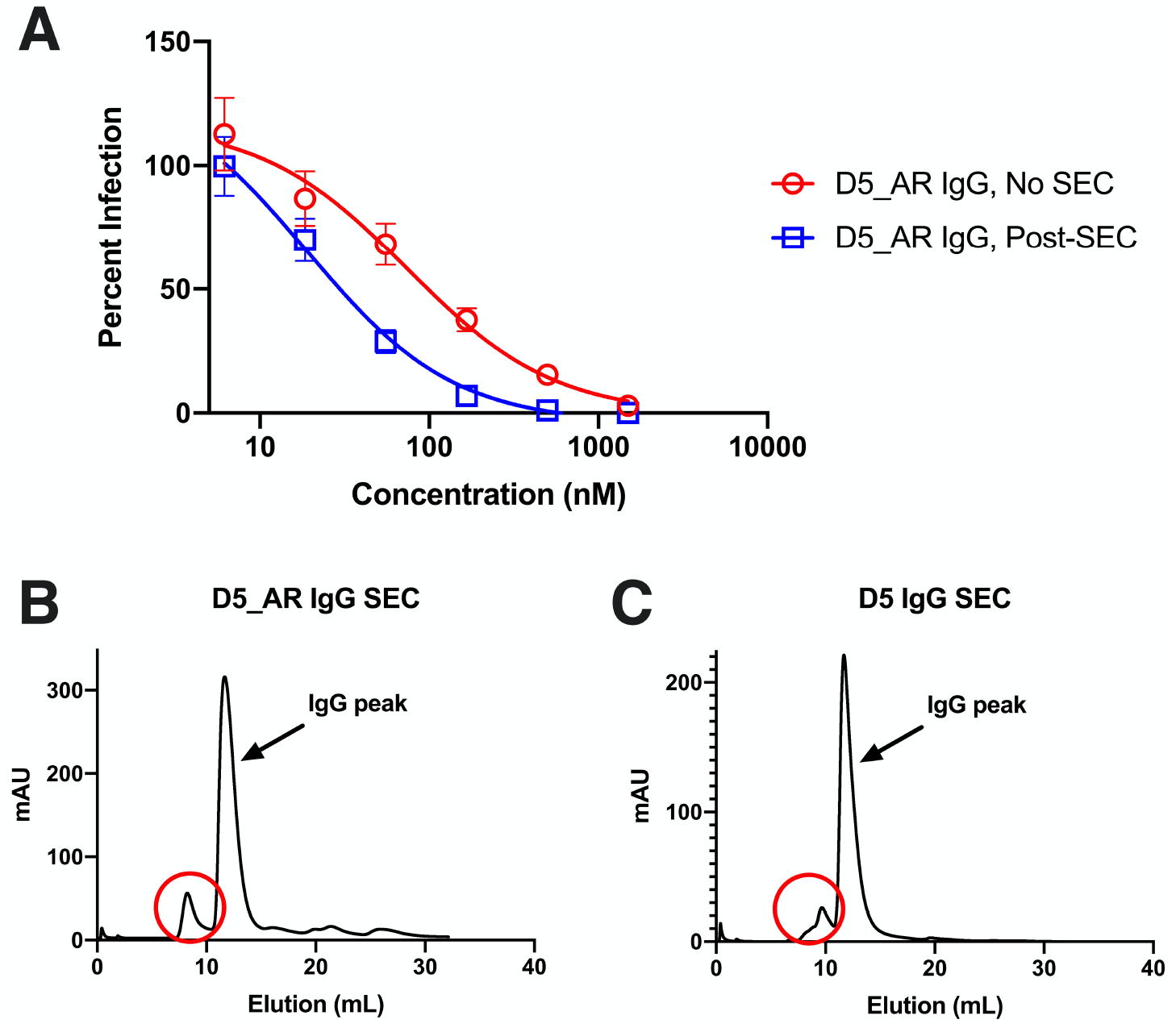
Purification via size exclusion chromatography (SEC) impacts the neutralization potency of D5_AR and reveals possible aggregation. **(A)** After purification by gel filtration, D5_AR IgG preparations demonstrated enhanced neutralization activity. Each data point represents the mean with standard error of mean (n=2). **(B and C)** UV traces from SEC reveal possible protein aggregates (circled in red) for both D5 and D5_AR IgG.

**Supplemental Table 1:**
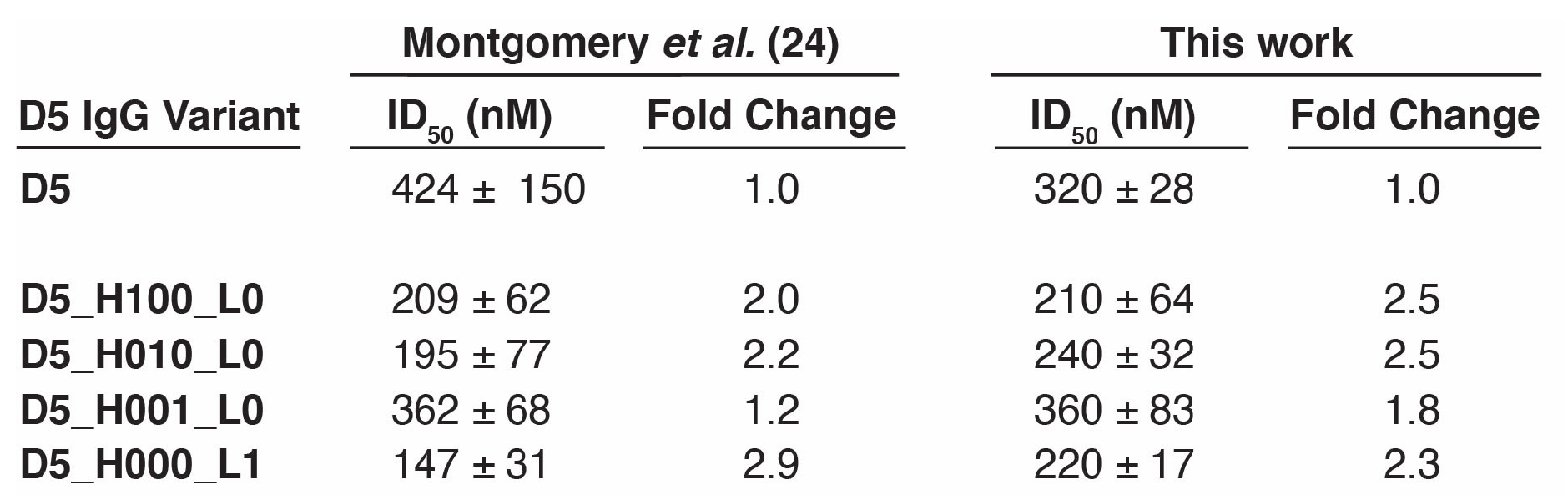
Single-CDR mutants neutralize pseudotyped HIV-1 HXB2 in a fashion consistent with the report of Montgomery *et al*. (24). The ID_50_ for this work is represented by the geometric mean and standard error of the mean of replicate experiments (n=2). For each infection assay, the fold enhancement versus D5 was calculated (ID_50, D5_ / ID_50, D5 variant_); reported fold enhancement is the geometric mean (n=2).

## Notes

### Competing Interest Statement

The authors have declared no competing interest.

